# Preserving condensate structure and composition by lowering sequence complexity

**DOI:** 10.1101/2023.11.29.569249

**Authors:** Amogh Sood, Bin Zhang

## Abstract

Biological condensates play a vital role in organizing cellular chemistry. They selectively partition biomolecules, preventing unwanted cross-talk and buffering against chemical noise. Intrinsically disordered proteins (IDPs) serve as primary components of these condensates due to their flexibility and ability to engage in multivalent, nonspecific interactions, leading to spontaneous aggregation. Theoretical advancements are critical at connecting IDP sequences with condensate emergent properties to establish the so-called molecular grammar. We proposed an extension to the stickers and spacers model, incorporating non-specific pairwise interactions between spacers alongside specific interactions among stickers. Our investigation revealed that while spacer interactions contribute to phase separation and co-condensation, their non-specific nature leads to disorganized condensates. Specific sticker-sticker interactions drive the formation of condensates with well-defined structures and molecular composition. We discussed how evolutionary pressures might emerge to affect these interactions, leading to the prevalence of low complexity domains in IDP sequences. These domains suppress spurious interactions and facilitate the formation of biologically meaningful condensates.

**Significance Statement:** Biomolecular condensates serve as pivotal mechanisms in cellular organization, often characterized by an abundance of intrinsically disordered proteins (IDPs) that undergo frequent mutations in their sequences. Despite this, IDP sequences exhibit non-random patterns, yet the precise relationship between these sequences and the emergent properties of condensates remains unclear. To address this gap, we propose a molecular theory that delineates how various sequence features of IDPs contribute to the organization and composition of condensates. This theory not only sheds light on the evolution of IDPs but also elucidates the emergence of non-random sequence patterns as essential elements for the formation of functional condensates. Correspondingly, we posit that the prevalence of low-complexity regions within IDPs is a result of evolutionary selection.

## Introduction

Biomolecular condensates serve as important mechanisms for the hierarchical and vectorial organization of chemistry within cells.^1–7^ They are non-stoichiometric assemblies of biomolecules that can form via spontaneous or driven processes and exhibit characteristics of phase separation and percolation.^8–10^ The chemical environment and physicochemical conditions within condensates are distinct from their surroundings,^11–18^ enabling cells to selectively partition biomolecules, prevent unwanted cross-talk and interference between various biochemical pathways, and buffer against chemical noise.

In recent years, these condensates have gained significant prominence and attracted substantial research interest. They have been found to primarily form through the involvement of intrinsically disordered proteins (IDPs).^19–22^ These unique proteins lack a fixed threedimensional structure, enabling them to be highly flexible and engage in multivalent interactions. Multivalent interactions involve the binding of multiple partners simultaneously and are often nonspecific in nature, leading to the spontaneous aggregation and condensation of various cellular components. Many research groups have attempted to establish what is known as the “molecular grammar”,^23–34^ which connects amino acid sequences with protein phase behaviors and the collective physical properties of condensates.

Establishing a theoretical framework is crucial for advancing our understanding of biological condensates. The “stickers and spacers” model, originally developed within the context of polymer gelation theory,^35–37^ has gained popularity for modeling protein condensates.^1,2,25,38–40^ In this model, biomolecules within condensates are envisioned as possessing two distinct functional components: “stickers” and “spacers”. Stickers represent specific molecular domains or motifs with a high affinity for one another, facilitating interaction and bringing molecules into close proximity, thereby contributing to the condensate’s cohesive and ordered structure. Conversely, spacers act as flexible linkers connecting the stickers, enabling the dynamic and transient nature of interactions within the condensate. This model offers a conceptual framework that assists in interpreting experimental observations regarding condensate stability and material properties.^41^

The simplicity of the stickers and spacers model, crucial for its theoretical elegance, encounters limitations when applied to realistic biomolecules. Specifically, when dealing with an IDP, the identification of stickers and spacers is often challenging.^1,3,42,43^ Most amino acids undergo some level of interactions, and determining when to disregard these interactions to categorize them as spacers remains ambiguous. We propose that instead of entirely disregarding their interactions, it’s essential to consider the interactions between spacer sequences and explore their impact on the physical properties of condensates. This exploration could offer insights into the evolutionary pressure of IDP sequences, arising from the interplay between sticker and spacer sequences. Moreover, such an investigation could enhance our understanding of real-world biological condensates, where spacers might deviate from ideal behavior but still participate in weak interactions.

We expanded the stickers and spacers model to include explicit non-specific pairwise interactions between spacers, alongside the specific interactions among sticker motifs. This extension allowed us to systematically explore the intricate interplay between specific and non-specific interactions in determining the structural and compositional properties of condensates. Our investigation revealed that spacer interactions contribute to phase separation and the co-condensation of multiple molecules. However, the non-specific nature of these interactions results in disorganized condensates with undefined molecular compositions. In contrast, specific sticker-sticker interactions drive the formation of condensates with robust contacts and precise compositions. Subsequently, we discussed the implications of our theory for the evolution of protein sequences, asserting the existence of evolutionary constraints even on segments of protein sequences that interact non-specifically. These constraints ensure the functionality of condensates. This evolutionary pressure naturally favors the emergence of low complexity domains to suppress spurious interactions, facilitating the formation of biologically meaningful condensates.

## Theory

### Stickers and random spacers model

We present a generalized version of the stickers and spacers model to investigate the phase behaviors of associative polymers (Fig. 1). These polymers consist of *N* monomers, with *f* privileged monomers that exhibit specific attractive interactions, denoted by a strength of −*u*_*a*_, leading to the formation of noncovalent, physical bonds. We refer to these privileged monomers as stickers. For simplicity, we consider stickers composed of a single monomer. However, it’s important to note that in biological sequences, stickers may consist of multiple amino acids. The remaining monomers are designated as spacers.

**Figure 1:**
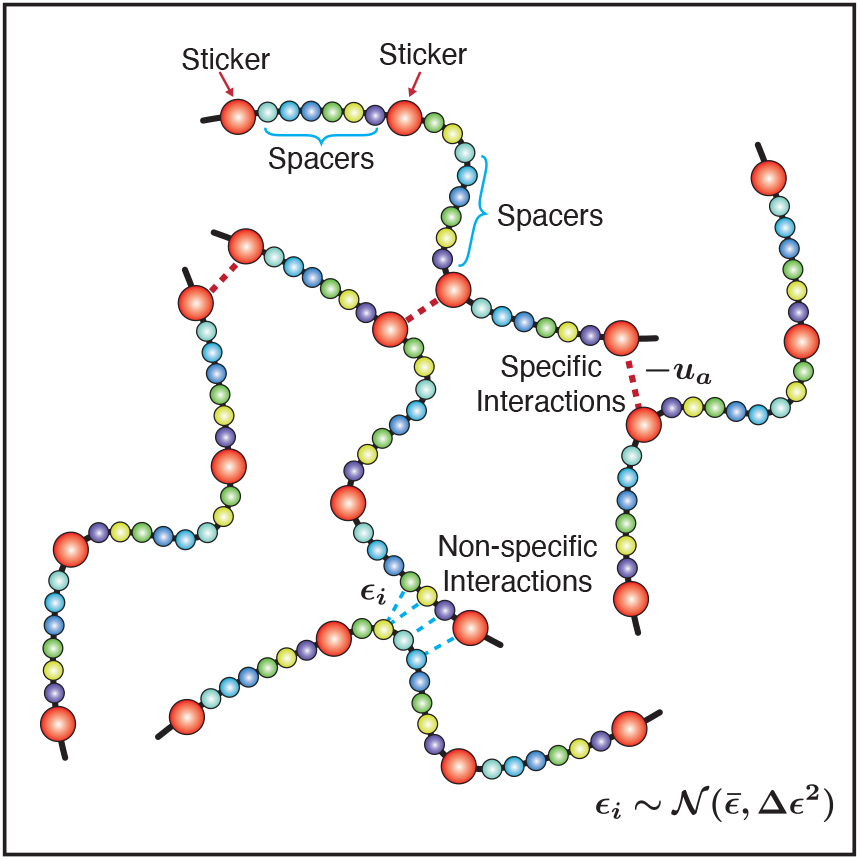
A schematic illustration of the stickers and random spacers model. Red spheres indicate stickers that interact specifically, and the strength for sticker-sticker interactions is a well-defined number, *−u*_*a*_. We indicate the random spacers using shades of blue-green. These contribute non-specific interactions, and the pairwise interactions between adjacent pairs of spacers are chosen from a normal distribution, 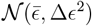.

In a departure from the traditional model, we consider the interaction energy between a pair of spacers, or a spacer-sticker pair, to be a random variable, *ϵ*_*i*_, drawn from a normal distribution, 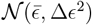, with mean 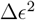 and variance Δ*ϵ*^2^. The contribution to the total energy for a given configuration on the lattice from non-specific interactions is

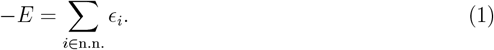

The summation denotes a sum over neighboring spacer-spacer pairs and non-bonded sticker and spacer pairs. Our use of random energy follows the tradition of protein folding theory,^44,45^ allowing the derivation of expressions with a mean field theory^46^ that are not specific to considerations of any particular sequence.

In the following, we explore two systems to investigate the influence of spacer interactions on the organization and compositions of condensates. Initially, we examine a homotypic system consisting of a single polymeric species encompassing both specifically interacting sticker moieties and non-specifically interacting spacer residues. In this homotypic system, we formulate a mean-field free energy and investigate percolation coupled with phase separation. Our analysis reveals how the distribution of non-specific spacer interactions impacts the critical temperature, critical concentration, gel point, and degree of conversion. Subsequently, we extend this model to a heterotypic system involving two polymeric species, denoted as *A* and *B*. We highlight the necessity of finely tuning non-specific interactions to ensure robust composition in the dense phase for *A* − *B* mixtures.

### Phase behaviors of a single component system

We begin by examining a single-component system comprising *n*_*p*_ identical polymer chains on a lattice containing *n* sites. Let *φ ≡ n*_*p*_*N/n* represent the fraction of sites occupied by the polymer. The formation of *n*_*p*_*m* bonds occurs between 2*n*_*p*_*m* stickers, and we define the degree of conversion as 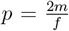. Here, the *f* sticker segments are assumed to be uniformly distributed, effectively partitioning the chain into *f* + 1 segments. We denote the expected length of each individual segment as *l*. By symmetry, (*f* + 1)*l* = *N* − *f* . Consequently, 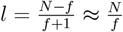 for *N ≫ f* and *f ≫* 1.

The partition function for forming *n*_*p*_*m* bonds between 2*n*_*p*_*m* stickers is,

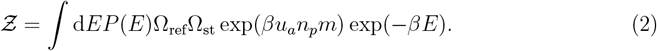

Here, the probability distribution of the energy, *P* (*E*), is Gaussian, since *E* is a sum of independent random variables following the Gaussian distribution (Eq. 1). The mean and variance of *P* (*E*) are given by

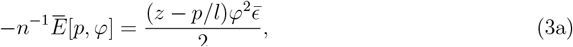

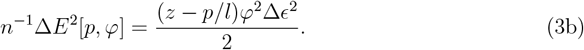

Here, *z* denotes the coordination number of the lattice. The term *p/l* appears to account for the fact that once a sticker forms a physical bond with another sticker, it is no longer available to interact with spacers. *β* = (*k*_*B*_*T* )^*−*1^ where *k*_*B*_ is the Boltzmann’s constant. For convenience, we set *k*_*B*_ = 1 and report all parameters in natural units.

Ω_ref_ is the number of ways of placing the *n*_*p*_ polymers on the and is a standard result in polymer physics.^47^ Its expression is given by,

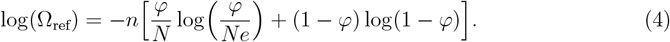

Ω_st_ corresponds to configurational entropy of forming sticker-sticker bonds. Following Semenov and Rubinstein ^35^, we derive the expression as

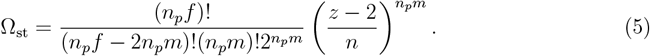

In deriving the above expression, we assumed that the chains are strongly overlapping (*φ >> φ*_overlap_ *∼ N*^*−*1*/*2^) and inter-chain bonds between stickers dominate over intra-chain ones. Thus, to arrive at Ω_st_, we first count the number of ways of choosing 2*n*_*p*_*m* stickers out of *n*_*p*_*f* stickers to form bonds. We then multiply this number by the number of ways of pairing 2*n*_*p*_*m* stickers together which is given by (2*n*_*p*_*m −* 1)!!. Finally, we multiply this by the probability that all of the chosen stickers are neighboring each other. We also used the relation 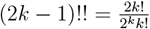 to further simplify the expression finally yielding Eq. 5.

The integration in Eq. 2 is readily performed, and we use *−βF* = log(*Ƶ*) to obtain the free energy. The free energy must be minimised with respect to the degree of conversion, *p*, yielding the condition

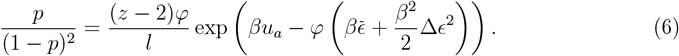

Remembering that *p* ∈ [0, 1], we derive

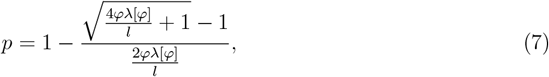

where 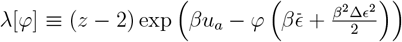. We substitute the condition, Eq. 6, into the original expression for *βF*, to obtain the free energy density 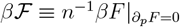 as

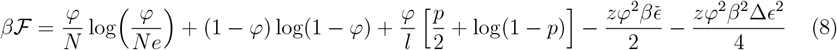

Eq. 8 provides the starting point for deriving equilibrium properties of the system. For example, to determine the critical behaviour we compute the chemical potential, *µ* = *∂*_*φ*_*ℱ*. The critical point is then the intersection of the nullclines *∂*_*φ*_*µ* = 0 and 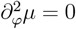. Additionally, we also consider the concentration of chains with all *f* stickers free, *𝒞*_free_, defined as,

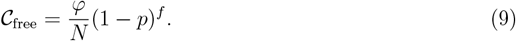

The gel line is obtained by the condition *∂*_*φ*_*𝒞*_free_ = 0. We also compute *φ*_dense_, the concentration in the dense phase, by using Π *≈* 0, where Π is the osmotic pressure given by the relation,

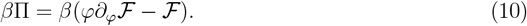

The concentration in the dilute phase (*φ*_dilute_) can then be obtained by equating the the chemical potentials in the two phases.

### Phase behaviors for a two component system

To further understand the influence of spacer interactions on condensate composition, we examine systems composed of two types of chains, denoted as *A* and *B*. Here, *n*_*a*_ and *n*_*b*_ represent the number of chains, while *N*_*a*_ and *N*_*b*_ denote the degree of polymerization for chains *A* and *B*, respectively. Chain *A* comprises *f*_*a*_ stickers, whereas chain *B* consists of *f*_*b*_ stickers. We permit specific *A − B* sticker-sticker interactions (*−u*_*ab*_) while prohibiting self-interactions (i.e., *u*_*aa*_ = *u*_*bb*_ = 0). This assumption mirrors a common scenario involving two proteins with specific binding.

The partition function for the two component system takes the familiar form,

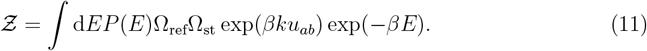

We once again assume the spacer-spacer and spacer-sticker interactions between *A − A, B − B* and *A − B* chains are each drawn from a normal distribution 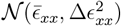 where, *xx* ∈ *{aa, bb, ab}*. Correspondingly, *P* (*E*) is Gaussian with mean and variance given by

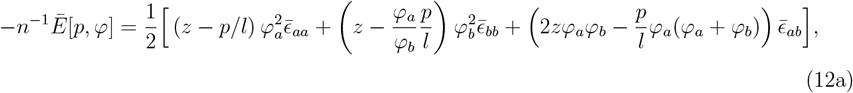

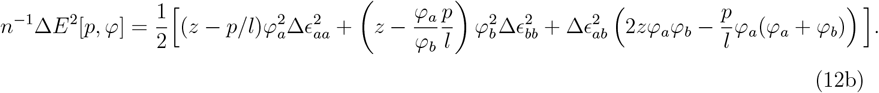

The mixing entropy is given by,^47^

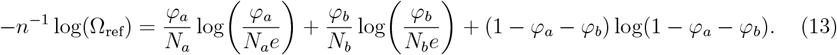

We assume the existence of *k* pairs of *A−B* sticker-sticker bonds in the system. Therefore, 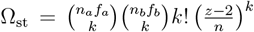. In order to define the degree of conversion we note that the maximum possible number of bonds is min*{n*_*a*_*f*_*a*_, *n*_*b*_*f*_*b*_*}*, and without loss of generality we can assume it is chain *A*.^39^ Therefore, we define *p*_*ab*_ *≡ k/n*_*a*_*f*_*a*_. For algebraic convenience, we assume 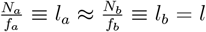. Consequently, the configurational entropy of the sticker-sticker bonds can be readily computed as:

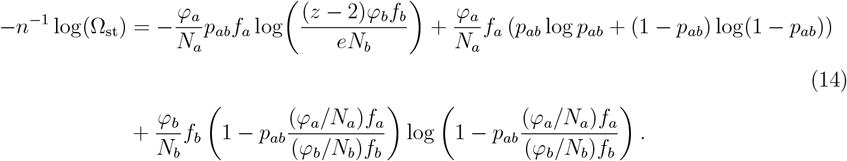

Here we have used the Stirling’s approximation (i.e, for *n >>* 1, log(*n*!) *≈ n* log *n − n* + *O*(log *n*)) and the following helpful identities, 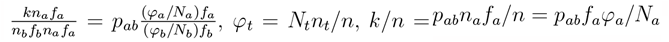, *k/n* = *p*_*ab*_*n*_*a*_*f*_*a*_*/n* = *p*_*ab*_*f*_*a*_*φ*_*a*_*/N*_*a*_.

We perform the integration in Eq. 11 and use *−βF* = log(Ƶ) to obtain the free energy. We then minimise it with respect to the degree of conversion, *p*_*ab*_, to yield,

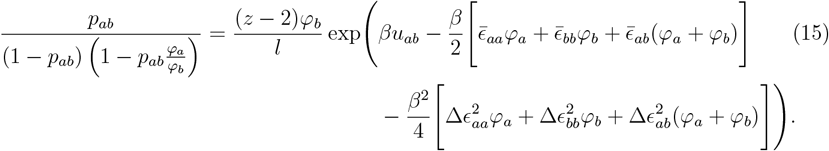

We substitute the above expression into the original expression for *βF* to obtain the free energy density 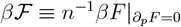

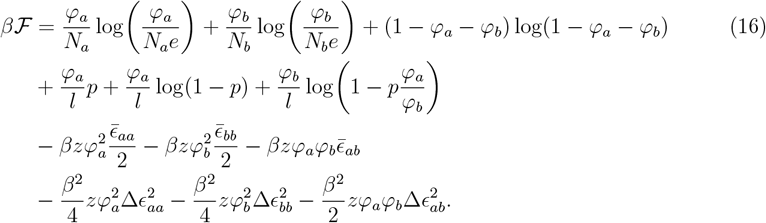

One can recover the expressions derived for the homotypic case from the above equation by replacing *φ*_*a*_ = *φ*_*b*_ *≡ φ*, up to a *φ →* 2*φ* transformation.

For the two component system, the spinodal can be obtained from the nullcline of the determinant of the hessian matrix of *ℱ*, namely, det(ℋ_*F*_ ) = 0. We determine the bimodal by solving the following system of equations

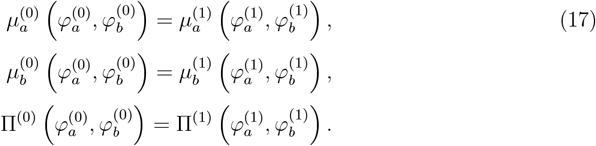

where, 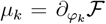 and 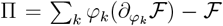. These equations ensure that the chemical potentials and the osmotic pressure are identical between coexisting phases. We solve the above equations by numerically finding roots to the the function 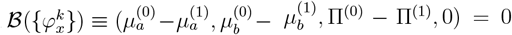 using standard python libraries. More details on the numerical solutions are provided in the Supporting Information.

## Results

### Non-specific spacer interactions facilitate phase separation

As described in the Theory section, we introduce a novel model designed to investigate the phase behavior of condensate-forming proteins (Fig. 1). Expanding upon the stickers and spacers model,^1,2,35–37^ we identify specific chemical groups within proteins that exhibit robust interactions as stickers. However, contrary to prevalent approaches in contemporary literature, we account for heterogeneity in interactions among spacers. For simplicity, we assume that the strength of spacer interactions follows a normal distribution, 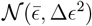, characterized by a mean of 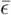 and a variance of Δ*ϵ*^2^. This variability accommodates the diversity of amino acids, resulting in weak yet significant interactions.^44,48,49^ We refer to this model as the “stickers and random spacers” model, or STARS.

We compute the complete phase diagram of the STARS model to investigate the impact of nonspecific spacer interactions on condensate behaviors. For a protein solution with identical molecules, we derive its free energy density *ℱ* (Eq. 8). From this, we calculate the chemical potentials as *µ* = *∂*_*φ*_*ℱ*, where *φ* represents the polymer volume fraction. The binodal is obtained by equating the chemical potentials of each component in coexisting phases, while the spinodal is obtained as the nullcline of *∂*_*φ*_*µ* = 0. These lines delineate the boundaries among the unstable, meta-stable, and stable regions of the phase diagram. The critical point (*T*_*c*_, *φ*_*c*_) denotes the temperature and concentration at which the solution first becomes unstable, aiding our understanding of the macromolecular solubility of the system.

We present the phase diagram for the STARS model in Fig. 2a. The sticker interactions are set as *u*_*a*_ = 5*k*_*B*_*T* and Δ*ϵ* = 2*k*_*B*_*T* with 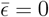. Additional model parameters are included in the caption. For comparison, we compute the phase diagram for a stickers and spacers model with similar parameters (Fig. 2b). Similar to the stickers and spacers model,^50,51^ the STARS model exhibits both a percolation (gelation) transition and phase separation. At high temperatures with *T > T*_*c*_, the system can undergo a sol-gel transition without phase separation. Decreasing the temperature further leads to phase separation coupled with gelation.

**Figure 2:**
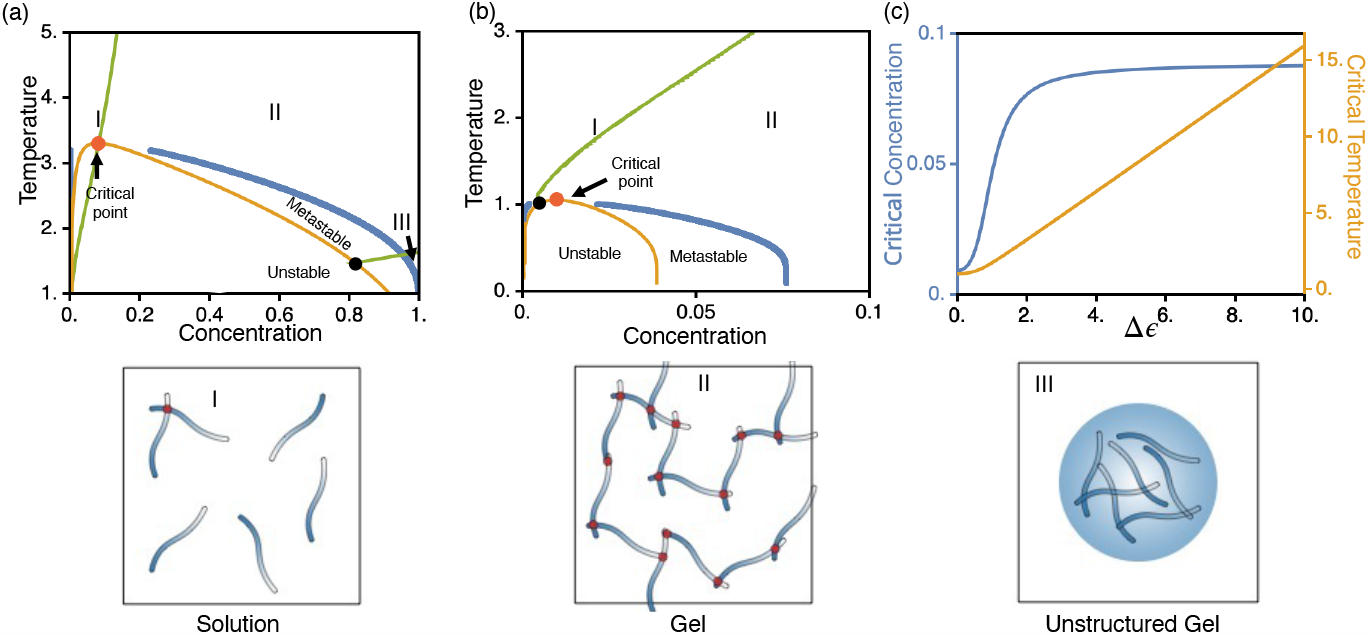
Phase behavior of the stickers and random spacers model. (a, b) Phase diagrams for the STARS model with Δ*ϵ* = 2 (a) and the stickers and spacers model with Δ*ϵ* = 0 (b). We plot the spinodal (orange) and binodal (blue) curves that demarcate the boundaries between the stable, meta-stable and unstable regions in the phase diagram. The critical point is highlighted in red. For the STARS model, the gel line (green) crosses the binodal twice, partitioning the stable phase into three regions. Illustrative configurations for the three regions corresponding to the solution phase, the gel phase, and the unstructured gel are shown in the bottom. (c) Dependence of the critical point on the strength of non-specific interaction among spacers, Δ*ϵ*. We set *u*_*a*_ = 5, 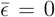, *l* = 10, *N* = 100, and *z* = 6 when computing the phase diagrams.

The inclusion of spacer interactions also leads to quantitative alterations in the phase diagram. In Fig. 2c, we observe a sharp rise in the critical temperature *T*_*c*_ with increasing Δ*ϵ*, thereby promoting phase separation. The critical concentration, *φ*_*c*_, follows a similar trend, albeit exhibiting a slight increase at larger values of Δ*ϵ* before quickly plateauing. Notably, these trends remain consistent regardless of other system parameters (Figs. S1 and S2). Spacer interactions, while chosen from a normal distribution with a zero mean, can produce contacts with negative energies that are favored by the Boltzmann factor, as defined in the partition function (Eq. 2). Consequently, these contacts stabilize the condensed phase, thereby facilitating phase separation.

### Non-specific spacer interactions modulate condensate organization

Fig. 2c indicates that diversifying IDP sequences might increase the variation in their interaction energies, potentially contributing positively to phase separation. However, it seems counter-intuitive that many IDPs, known to participate in condensate formation, would evolve sequences featuring low complexity regions.^52–55^ These regions often contain amino acid repeats with suppressed sequence diversification and interaction patterns, leading to smaller Δ*ϵ* values.

We propose that spacer-spacer interactions could negatively impact condensate structures arising from physical crosslinks between stickers. These crosslinks differentiate biomolecular condensates from simple liquids, generating heterogeneous environments with specific protein-protein interfaces. Robust contacts inside condensates could facilitate fast processing of intermediates, such as in metabolic channeling.^56^ Additionally, physical crosslinks influence the viscoelastic and rheological properties crucial for condensate function.^1,2,50^ For instance, differences in viscoelasticity between nucleolar core and outer layers strongly affect ribosomal assembly.^57^

To characterize condensate structure, we introduce the degree of conversion, *p*, measuring the fraction of bonded stickers. Additionally, we study the dependence of *p* on Δ*ϵ* at a constant temperature *T* = 1. For the parameters outlined in the caption of Fig. 3, the system undergoes phase separation coupled with gelation at this temperature. We denote the polymer volume fraction of the dense phase as *φ*_dense_ and use it to determine *p* from Eq. 7. As depicted in Fig. 3a, the degree of conversion initially increases slightly, reaching a maximum before decreasing to zero with increasing Δ*ϵ*. This suggests that spacer-spacer interactions not only influence the overall macromolecular solubility but also mediate network properties of the stickers.

**Figure 3:**
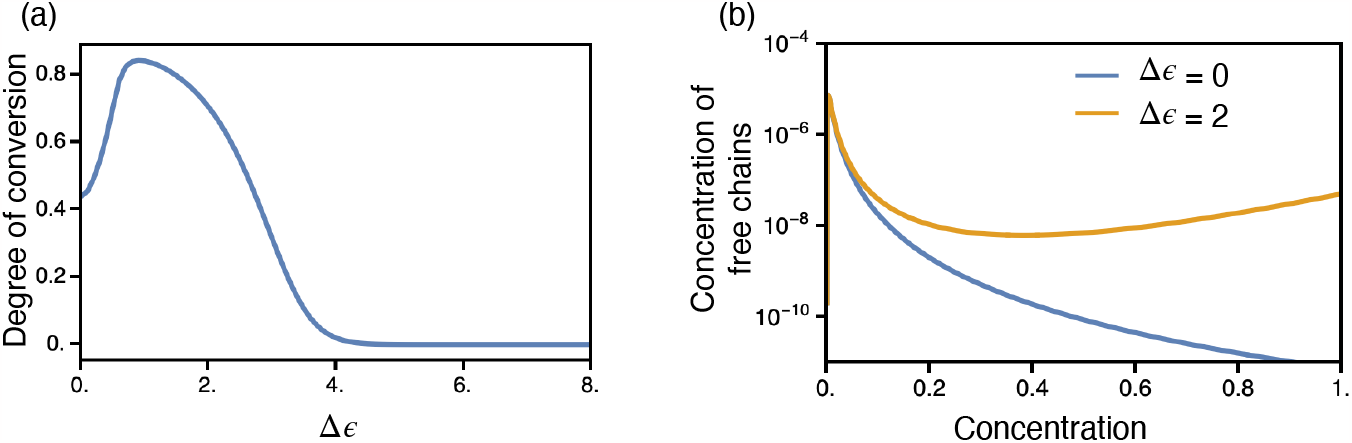
Impact of non-specific interactions among spacer interactions on the network properties of condensates. (a) The degree of conversion evaluated at *φ*_dense_ (the concentration of polymers in the dense phase) shows a moderate increase followed by a subsequent decrease as we widen the spread of the spacer-spacer interaction energy distribution by increasing Δ*ϵ*.(b)The concentration of free chains (with all stickers free) increases with concentration in the pre-gel regime and reaches a maximum at the gel-point. It monotonically decreases in the post-gel regime when Δ*ϵ* = 0, but exhibits non-monotonic behavior for Δ*ϵ≠* 0. We set *u*_*a*_ = 5, *T* = 1, 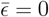, *l* = 10, *N* = 100, and *z* = 6.

Moreover, spacer-spacer interactions can qualitatively change the phase diagram. We study *𝒞*_free_, measuring the concentration of polymer chains with all stickers free (i.e., not bound to other stickers, Eq. 9). As per established theories, for Δ*ϵ* = 0, *𝒞*_free_ first increases with concentration (*φ*) in the pre-gel regime, achieving a maximum at the gel point (Fig. 3b, blue). Subsequently, in the post-gel regime, *𝒞*_free_ monotonically decreases to zero as an increasing number of stickers form crosslinks.^35^ Consequently, the maximum of *𝒞*_free_ often defines the gel line (*∂*_*φ*_*𝒞*_free_ = 0), depicted in Fig. 2 as the green curves.

Strikingly, we note that the monotonic decrease in *𝒞*_free_ as a function of increasing *φ* in the post-gel regime for Δ*ϵ*≠ 0 no longer holds true. As shown by the orange line in Fig. 3b, *𝒞*_free_ decreases to a minimum before increasing again at higher concentrations. Once again, these observations remain qualitatively insensitive to the exact choice of system parameters (Figs S1 and S2).

This non-monotonic behavior results in the first derivative of *𝒞*_free_ crossing the binodal twice in Fig. 2a, partitioning the stable regions into three. Regions I and II resemble the typical solution and gel phase found in the stickers and spacers model (see Fig. 2b). However, in region III, *𝒞*_free_ increases again due to a drop in sticker-sticker crosslinks. This transition leads to a less structured gel due to the competition between spacer interactions and the formation of sticker-sticker bonds.

### Non-specific spacer interactions modulate condensate composition

Until now, our focus has been on single-component systems. However, biomolecular condensates within cells commonly consist of multiple molecules. ^8,10,58–61^ These condensates possess well-defined compositions, allowing only certain molecules to selectively partition into them.^62^ Maintaining such a specific composition is crucial for their function in both physiological contexts and potential therapeutic applications.^13,63–65^ To explore further, we investigated whether spacer interactions influence condensate composition in a minimal system comprising two components.

We aimed to model a scenario where *A* serves as the host, incorporating a polymeric guest *B*. Consequently, we focused on homotypic *A* − *A* interactions (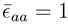, Δ*ϵ*_*aa*_ = 0) to facilitate the demixing of polymer *A* from *B*.^16^ To simplify, our attention was directed toward sequence diversification in protein *B*. This diversification results in both self-interactions and cross-interactions with *A*. Hence, we set both *ϵ*_*bb*_ and *ϵ*_*ab*_ as random variables with variance Δ*ϵ*_*bb*_ *≡* Δ*ϵ*_*ab*_ and zero-mean (for convenience).

For this two-component system, we derived the free energy expression (Eq. 16) to study its phase behaviors. Fig. 4 illustrates the phase behavior at constant temperature (*T* = 1) as a function of composition in the *φ*_*a*_ −*φ*_*b*_ plane, considering varying strengths of specific (*u*_*ab*_) and nonspecific interactions (Δ*ϵ*_*bb*_, Δ*ϵ*_*ab*_). The orange lines denote the spinodal, while the blue lines represent the binodal. Additionally, the gray lines represent tie lines connecting coexisting phases.

**Figure 4:**
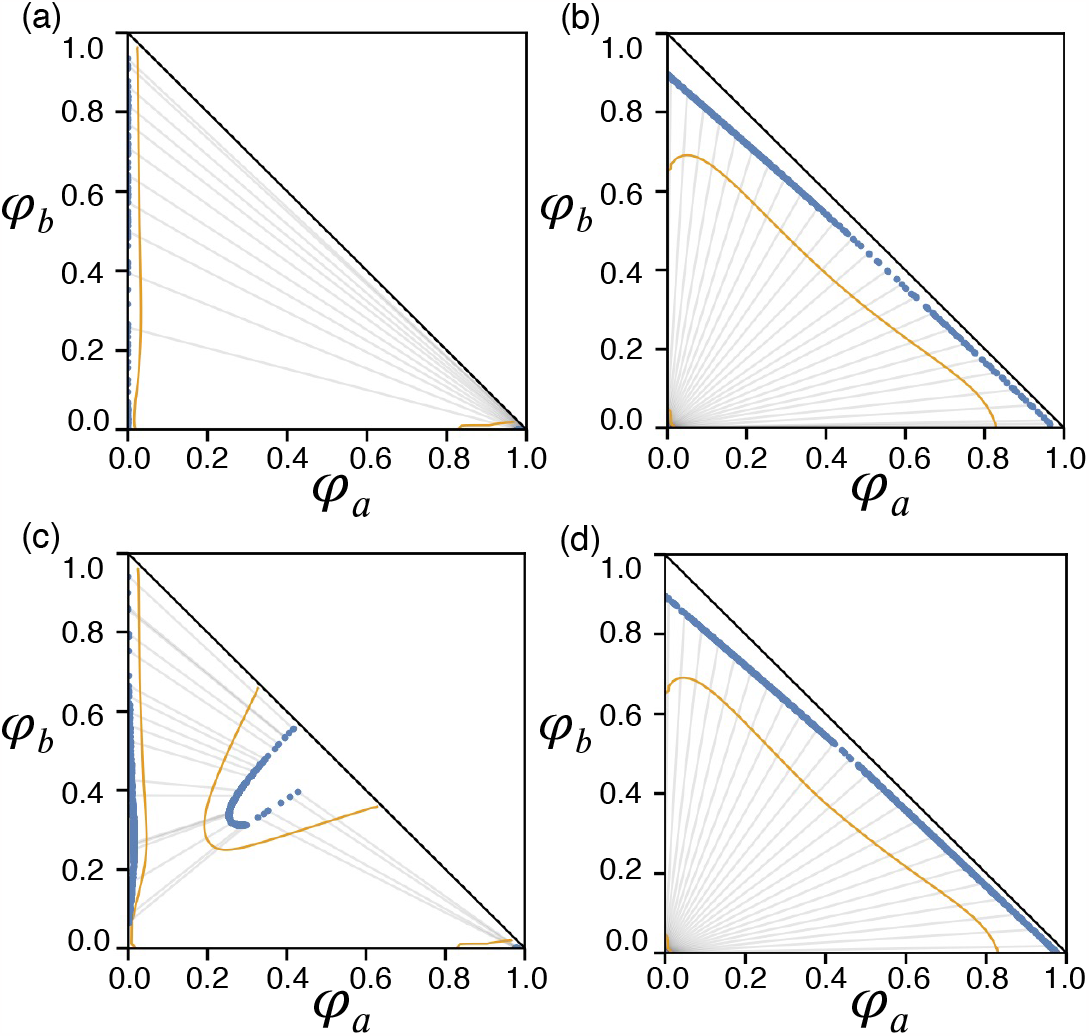
Phase diagrams showing the spinodal line (orange, solid) and binodal line (blue, dots) for a two component system with (a) *u*_*ab*_ = 2, Δ*ϵ*_*bb*_ = Δ*ϵ*_*ab*_ = 0, (b) *u*_*ab*_ = 2, Δ*ϵ*_*bb*_ = Δ*ϵ*_*ab*_ = 1.1, (c) *u*_*ab*_ = 5, Δ*ϵ*_*bb*_ = Δ*ϵ*_*ab*_ = 0, and (d) *u*_*ab*_ = 5, Δ*ϵ*_*bb*_ = Δ*ϵ*_*ab*_ = 1.1. The tie-lines (light grey, solid) connects co-existing points on the binodal. We set *N*_*a*_ = *N*_*b*_ = 10, *l* = 2, *z* = 6, 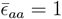, Δ*ϵ*_*aa*_ = 0, and 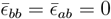 in all systems.

We observed that the inclusion of nonspecific spacer-spacer interactions facilitates the co-condensation of *A* and *B*. Specifically, in scenarios where sticker-sticker interactions are sufficiently weak (only 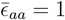 without any *A−B* or *B−B* spacer interactions), the negative slope of tie lines in Fig. 4a suggests a demixing behavior. Stable phases at the top left and bottom right exhibit enrichment in only one component without a balanced presence of both. However, upon introducing nonspecific spacer interactions in Fig. 4b, tie line slopes become positive, indicating stable phases enriched or depleted in both components simultaneously– signifying co-condensation. A similar behavior was observed by Deviri and Safran ^16^ using the Flory-Huggins theory. The observed splay in the tie lines in Fig. 4b indicates that slight concentration fluctuations in the dilute phase can lead to significantly distinct compositions in the dense phase.

Furthermore, enhancing interaction strength among stickers (while maintaining Δ*ϵ*_*ab*_ = Δ*ϵ*_*bb*_ = 0) also facilitates co-condensation. As shown in Fig. 4c, a new stable phase emerges within the diagram’s center. Tie lines connect this stable phase with regions depleted in one component at the top left and lower right corners. In this new stable region, *p*_*ab*_ *≈* 1. Additionally, the stable co-condensation phase has a narrower range of possible concentrations for *A* and *B*. This narrow concentration range is unlike the wider range supported in Fig. 4b and d, where a broader spectrum of mixing ratios exists in the stable condensed phase. In our current setup, the stable phase in Fig. 4c is centered around (0.5, 0.5) due to symmetric sticker distributions in *A* and *B*. Altering the interaction strengths among stickers or the relative abundance of *A* or *B* stickers can aid in promoting phases with varied compositions. A detailed survey of titrating Δ*ϵ*_*bb*_, Δ*ϵ*_*ab*_ on the phase diagram in both the weak (*u*_*ab*_ = 2) and strong (*u*_*ab*_ = 5) sticker-sticker interaction regimes is shown in the Supporting Information (Figs S3 and S4).

Thus, while both sticker-sticker and spacer interactions induce condensation of multiple components, sticker interactions robustly generate condensates with more defined molecular compositions and structures. This property can be advantageous in biological systems requiring precise rationing of different species to optimize efficiency in specific chemical processes. These observations remain qualitatively insensitive to the exact choice of system parameters (Figs S5 and S6).

## Conclusions and Discussion

We have introduced a new theory aimed at establishing connections between the emergent physical properties of condensates and protein sequences. Specifically, by extending from the stickers and spacers model, we investigate how the interplay between specific and nonspecific interactions can influence both the structural integrity and compositional specificity of biomolecular condensates. Specific interactions are defined as those exclusively formed between a pair of stickers due to their chemical selectivity, while non-specific interactions can occur among spacers and between spacers and non-bonded stickers. We analytically solved the new stickers and random spacers model to assess its phase behaviors.

Our findings demonstrate that non-specific spacer interactions have the capacity to promote phase separation and the condensation of multi-component systems. However, these non-specific interactions fail to generate condensates with robust organizational structures necessary for fine-tuning material properties and molecular compositions. Conversely, specific interactions show a similar ability to promote phase separation while establishing welldefined interaction networks within the condensate, resulting in precise compositions.

### Revisiting the definition of stickers and spacers

While it’s acknowledged that stickers and spacers are context-dependent, it’s common to identify potential stickers by pinpointing residues known to have the most pronounced impact on saturation concentrations.^3,66,67^ However, our study suggests that relying solely on critical behavior as a criterion for identifying potential stickers might be less robust. This is because spacers, characterized by non-specific interactions, can also influence both the critical and percolation behavior of the system. Alternatively, from a functional standpoint, we propose that stickers can also be identified as chemical groups that facilitate protein-protein interactions through specific binding. According to this definition, natural candidates for stickers are molecular recognition features (MoRFs) or short linear motifs (SLiMs). These consist of short stretches of adjacent amino acids essential for molecular recognition and protein binding.^68–70^ Regions undergoing significant mutations leading to higher levels of binding promiscuity, however, should be classified as spacers. Identifying stickers and spacers based on interaction specificity helps in understanding the sequence features of IDPs and their evolutionary conservation, as we discuss below.

### Evolutionary pressure and the rise of low complexity

Our study offers insight into the evolution of IDP sequences. IDPs are recognized for their high evolvability, characterized by rapid mutation rates.^71,72^ Unlike globular proteins that require well-defined 3D structures, IDPs theoretically possess a broader sequence space for exploration.^55^ However, IDPs do not explore all possible unfoldable sequences. Instead, they adopt simplified sequences enriched with low complexity domains, utilizing only a subset of amino acids. Furthermore, despite their poor conservation in alignments, recent studies have revealed that orthologous IDPs share many conserved molecular features,^73–75^ indicating non-randomness and suggesting evolutionary constraints that favor functionally fit sequence patterns.

Yet, the mere formation of biological condensates cannot explain the conserved features in IDP sequences. As discussed in the main text, the diversification of spacer sequences can effectively reduce saturation concentrations and facilitate phase separation. Therefore, to just optimize condensate stability, IDPs would not heavily favor low complexity domains that suppress interactions.

We propose that the formation of condensates with robust structures and compositions serves as the main pressure for IDP evolution. This pressure strengthens interactions among stickers while concurrently suppressing promiscuous interactions among spacers, leading to the prevalence of low complexity domains in IDPs. Lowering sequence complexity limits the range of possible amino acids within the sequence, naturally reducing the complexity and fluctuation of unwanted interactions. Additionally, for sequences of identical amino acid composition, interaction complexity can be further reduced by creating repeating, alternating aromatic residue sequences, such as FG-repeats in nucleoporins or YG-repeats in FUS-LCD.^3,76,77^ These alternating sequences are known to exhibit weaker interactions compared to sequences where amino acids of the same type are clustered together.^78^

The suggested evolutionary pressure inherently drives the preservation of stickers to maintain condensate compositional specificity. Molecular recognition features and short linear motifs, acknowledged for their evolutionary conservation, support this notion.^73^ Conversely, spacers encounter an evolutionary push to diminish their complexity. This pressure doesn’t favor any particular sequence, hence contributing to their high mutation rate. Nevertheless, mutations resulting in excessively strong interactions, even when localized within the low complexity domains (i.e., spacers), can disrupt normal function as well.^79,80^

## Acknowledgments

The authors thank A. Athreya for helpful conversations. This work was supported by the National Institutes of Health (Grant R35GM133580).

## Code and Data Availability

Data presented in this study is available upon reasonable request to the corresponding author.

