## Supporting Information for "Preserving condensate structure and composition by lowering sequence complexity"

#### Contents

**Interrogating the Effect of Parameters $u_a, l$ on the Single Component System S-2**

**Interrogating the Effect of $\Delta\epsilon_{ab}, \Delta\epsilon_{bb}$ on the Phase Behavior of the Two Component System S-3**

**Interrogating the effects of $N, l$ on the Phase Behavior of the Two Component System S-3**

**Details of Numerical Computations S-4**

Computing spinodal, binodal and critical points for the single component system . S-4

Computing spinodal and binodal for the two component system . . . . . . . . . . S-4

### Interrogating the Effect of Parameters $u_a, l$ on the Single Component System

We demonstrate that the qualitative trends discussed in the main text remain robust despite variations in sticker strength ($u_a$) and patterning ($l = N/f$). In Fig. S1 (a) and (b), we qualitatively replicate the increase in critical temperature and concentration highlighted in the main text for $u_a = 2.5$ and $u_a = 10$, respectively. Similarly, in Fig. S1 (c) and (d), we observe a similar qualitative dependence of the degree of conversion on $\Delta\epsilon$ for $u_a = 2.5$ and $u_a = 10$, respectively. Finally, Fig. S1 (e) and (f) demonstrate, akin to the observations in the main text, an increase in the concentration of chains with all stickers free, $\mathcal{C}_{\text{free}}$, up to a maximum value in the pre-gel regime but a monotonic decrease in the post-gel regime for $\Delta\epsilon = 0$. $\mathcal{C}_{\text{free}}$ exhibits non-monotonic behavior in the post-gel regime for $\Delta\epsilon \neq 0$ for both $u_a = 2.5$ and $u_a = 10$, respectively.

The parameter $l$ can also modulate the amount of specific interactions in the system, and our qualitative observations and conclusions in the main text remain independent of the specific choice of $l$. In Fig. S2 (a) and (b), we qualitatively reproduce the increase in critical temperature and concentration outlined in the main text for $l = 20$ and $l = 5$, respectively. Similarly, in Fig. S2 (c) and (d), we note a qualitatively similar dependence of the degree of conversion on $\Delta\epsilon$ for $l = 20$ and $l = 5$, respectively. Finally, Fig. S2 (e) and (f) demonstrate, similar to the main text, an increase in the concentration of chains with all stickers free up to a maximum value in the pre-gel regime. However, this concentration monotonically decreases in the post-gel regime for $\Delta\epsilon = 0$, while displaying non-monotonic behavior in the post-gel regime for $\Delta\epsilon \neq 0$ for both $l = 20$ and $l = 5$, respectively.

### Interrogating the Effect of $\Delta\epsilon_{ab}, \Delta\epsilon_{bb}$ on the Phase Behavior of the Two Component System

In this section, we discuss the changes in the phase behavior of the two-component system as we modulate $\Delta\epsilon_{ab} = \Delta\epsilon_{bb}$ in both the low-sticker strength ($u_{ab} = 2$) and high-sticker strength ($u_{ab} = 5$) regimes. Since computing binodals is often numerically challenging, we plot the spinodals to efficiently scan the parameter space and gain qualitative insights (see Fig. S3 and Fig. S4). In Fig. S3a-e, we observe the system's transition from the demixing regime to the co-condensation regimes, depicting phase behavior at intermediate values of $\Delta\epsilon_{ab} = \Delta\epsilon_{bb}$ in the regime where sticker-sticker interaction strengths are relatively weak ($u_{ab} = 2$). Conversely, in Fig. S4a-e, we observe the system's transition from the demixing regime to the co-condensation regimes, exhibiting phase behavior at intermediate values of $\Delta\epsilon_{ab} = \Delta\epsilon_{bb}$ in the regime where sticker-sticker interaction strengths are relatively strong ($u_{ab} = 5$).

We observe the $U$-shaped region at the center of the phase diagram in Fig. S4a, primarily stabilized by $A - B$ cross-links, which also emerges in the weak sticker regime, albeit at $\Delta\epsilon_{ab} = \Delta\epsilon_{bb} \neq 0$ (Fig. S3b). Notably, this $U$-shaped region, characterized by a narrow composition range for the A-B mixtures, appears only at intermediate values of $\Delta\epsilon_{ab} = \Delta\epsilon_{bb}$. As concluded in the main text, higher values of $\Delta\epsilon_{ab} = \Delta\epsilon_{bb}$–indicating stronger spacer interactions–eliminate this region, resulting in condensates with poorly defined compositions.

### Interrogating the effects of $N, l$ on the Phase Behavior of the Two Component System

In Fig. S5, we qualitatively reproduce phase diagrams similar to those discussed in Fig. 4 of the main text, but now for longer polymer chains, specifically $N_a = N_b = 100$. All other parameters are held constant: $u_{ab} = 5, l = 2, z = 6, \bar{\epsilon_{aa}} = 1$.

Additionally, we observe that the parameter $l = N_x/f_x$ can significantly impact the phase behavior as it relates to the relative abundance of stickers. Increasing $l$ from $l = 2$ to $l = 10$, for a bead system with $N_a = N_b = 100$, is equivalent to reducing the number of stickers by a fifth. Consequently, maintaining the interaction strength at $u_{ab} = 5$ while setting $l = 10$ places us within the weak sticker regime previously discussed in the main text (see Fig. S6a-b). However, we can still qualitatively reproduce the phase behavior at strong sticker interactions by simultaneously increasing the sticker-sticker interaction strength to $u_{ab} = 10$ (Fig. S6c-d). This adjustment compensates for the reduction in sticker abundance caused by the increase in $l$, allowing us to maintain similar qualitative phase characteristics.

### Details of Numerical Computations

#### Computing spinodal, binodal and critical points for the single component system

The spinodal is given by the nullcline $\partial_\varphi \mu = 0$. The critical point is then the intersection of the nullclines $\partial_\varphi \mu = 0$ and $\partial^2_\varphi \mu = 0$.[S1] $\varphi_{\text{dense}}$ was estimated $\Pi \approx 0$.[S2] This was used as an initial guess in the root finding algorithm. Initialisation for $\varphi_{\text{dilute}}$ values for the root-finding algorithm were obtained using a line search in the interval $0 \leq \varphi_{\text{dilute}} < \varphi_{\text{dense}}$. Using these initialisations, the binodal was computed by equating the chemical potentials in the dilute and dense phases using a root-finding algorithm. We used *Mathematica* (Version 13.2) and the `FindRoot` function therein to execute these computations.

#### Computing spinodal and binodal for the two component system

For the two component system, the spinodal can be obtained from the nullcline of the determinant of the hessian matrix of $\mathcal{F}$, namely, $\det(\mathcal{H}_\mathcal{F}) = 0$. We determine the binodal by solving the following system of equations

$$\begin{aligned}
\mu_a^{(0)}\left(\varphi_a^{(0)},\varphi_b^{(0)}\right) &= \mu_a^{(1)}\left(\varphi_a^{(1)},\varphi_b^{(1)}\right), \\
\mu_b^{(0)}\left(\varphi_a^{(0)},\varphi_b^{(0)}\right) &= \mu_b^{(1)}\left(\varphi_a^{(1)},\varphi_b^{(1)}\right), \\
\Pi^{(0)}\left(\varphi_a^{(0)},\varphi_b^{(0)}\right) &= \Pi^{(1)}\left(\varphi_a^{(1)},\varphi_b^{(1)}\right).
\end{aligned} \tag{S1}$$

where, $\mu_k = \partial_{\varphi_k}\mathcal{F}$ and $\Pi = \sum_k \varphi_k(\partial_{\varphi_k}\mathcal{F}) - \mathcal{F}$. These equations ensure that the chemical potentials and the osmotic pressure are identical between coexisting phases. We solve the above equations by numerically finding roots to the the function $\mathcal{B}(\{\varphi_x^k\}) \equiv (\mu_a^{(0)} - \mu_a^{(1)}, \mu_b^{(0)} - \mu_b^{(1)}, \Pi^{(0)} - \Pi^{(1)}, 0) = 0$ using the *Scipy* library (Version 1.10.0) `scipy.optimize.root` function using the default implementation of Powell's hybrid method.[S3–S5] In addition, analytic derivatives and Jacobian's computed using the python *Sympy* library (Version 1.11.1) and *Numpy* (Version 1.24.1) library were utilised for other incidental numerical manipulations. Initial values for $\varphi_x^{(k)}$ for $x \in a, b$ provided to the root-finding algorithm, were chosen on a fine-grid in the triangle bounded $\varphi_a + \varphi_b \leq 1$ and $\varphi_a \geq 0$ and $\varphi_b \geq 0$. In general, precisely computing binodals can be numerically challenging, and we direct interested readers to sophisticated numerical approaches in the literature[S6–S12].

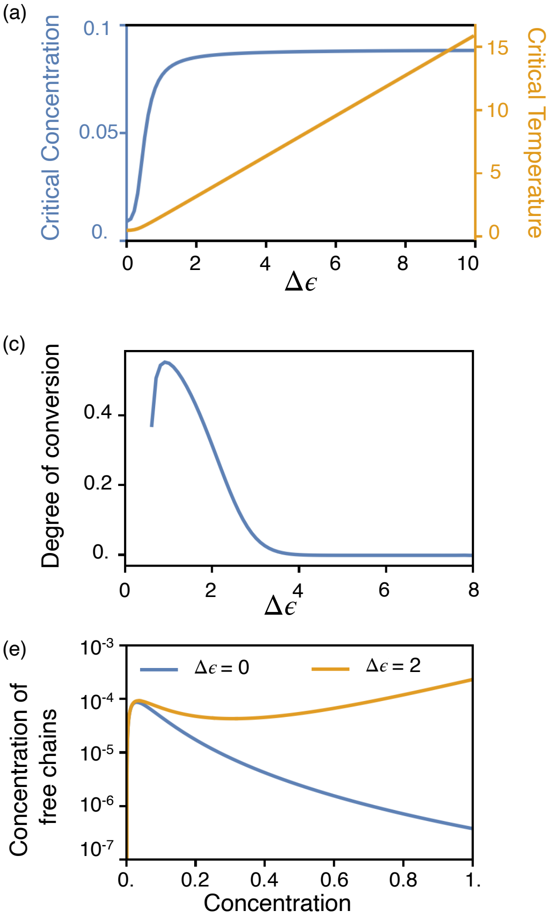


Figure S1: The results discussed in the main text for the one component system can be recapitulated for $u_a = 2.5$ and $u_a = 10$. We plot how the critical point shifts as we vary $\Delta\epsilon$ for (a) $u_a = 2.5$ and (b) $u_a = 10$. We also plot the degree of conversion evaluated at $\varphi_{\text{dense}}$ (concentration of polymers in the dense phase in the gel regime) as a function $\Delta\epsilon$ for (c) $u_a = 2.5$ and (d) $u_a = 10$ respectively. The concentration of free chains (with all stickers free) increases with concentration in the pre-gel regime and admits a maximum at the gel-point. The concentration of free chains is monotonically decreasing in the post-gel regime when $\Delta\epsilon = 0$ but exhibits non-monotonic behaviour for $\Delta\epsilon \neq 0$ for both (e) $u_a = 2.5$ and (f) $u_a = 10$ respectively. We set $\bar{\epsilon} = 0$, $l = 10$, $N = 100$, and $z = 6$. Additionaly, for (c)-(f) we held fixed $T = 1$. See text *Section: Interrogating the effect of parameters* $u_a, l$ *on the single component system* for more details.

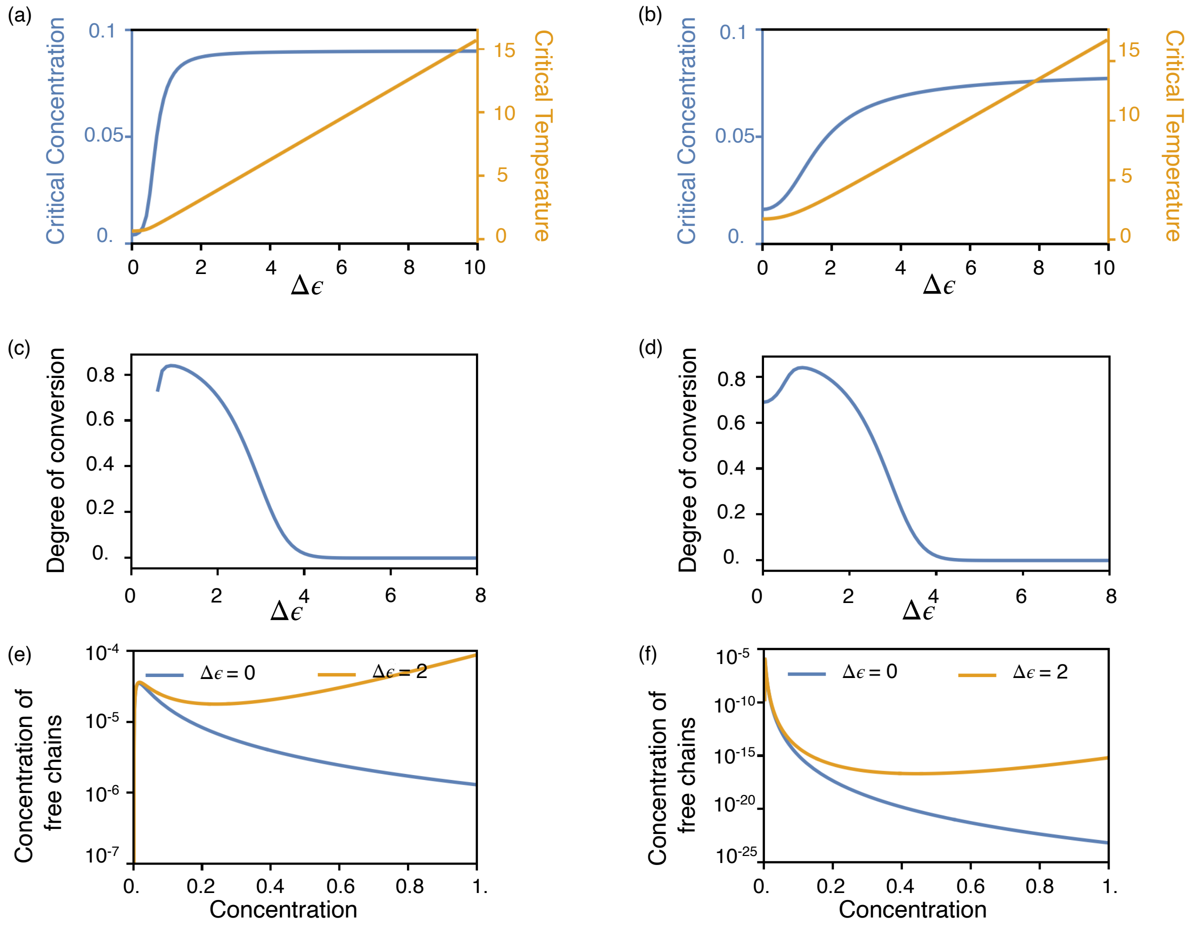


Figure S2: The results discussed in the main text for the one component system can be recapitulated for $l = 20$ and $l = 5$. We plot how the critical point shifts as we vary $\Delta\epsilon$ for (a) $l = 20$ and (b) $l = 5$. We also plot the degree of conversion evaluated at $\varphi_{\text{dense}}$ (concentration of polymers in the dense phase in the gel regime) as a function of $\Delta\epsilon$ for (c) $l = 20$ and (d) $l = 5$ respectively. The concentration of free chains (with all stickers free) increases with concentration in the pre-gel regime and admits a maximum at the gel-point. The concentration of free chains is monotonically decreasing in the post-gel regime when $\Delta\epsilon = 0$ but exhibits non-monotonic behaviour for $\Delta\epsilon \neq 0$ for both (e) $l = 20$ and (f) $l = 5$ respectively. We set $\bar{\epsilon} = 0$, $u_a = 5$, $N = 100$, and $z = 6$. Additionally, for (c)-(f) we held fixed $T = 1$. See text *Section: Interrogating the effect of parameters* $u_a, l$ *on the single component system* for more details.

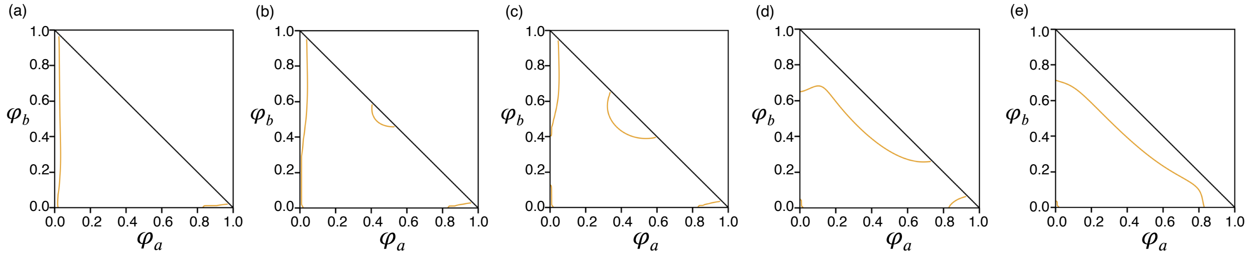


Figure S3: Phase diagrams showing the spinodal line (orange, solid) for a two component system with with $u_{ab} = 2$ (a) $\Delta\epsilon_{bb} = \Delta\epsilon_{ab} = 0$, (b) $\Delta\epsilon_{bb} = \Delta\epsilon_{ab} = 0.7$, (c) $\Delta\epsilon_{bb} = \Delta\epsilon_{ab} = 0.8$, (d) $\Delta\epsilon_{bb} = \Delta\epsilon_{ab} = 1.0$ and (e) $\Delta\epsilon_{bb} = \Delta\epsilon_{ab} = 1.1$ . We set $N_a = N_b = 10, l = 2, z = 6, \bar{\epsilon}_{aa} = 1, \Delta\epsilon_{aa} = 0$, and $\bar{\epsilon}_{bb} = \bar{\epsilon}_{ab} = 0$ in all systems. See text *Section: Interrogating the effects of* $\Delta\epsilon_{ab}, \Delta\epsilon_{bb}$ *on the Phase Behavior of the Two Component System* for more details.

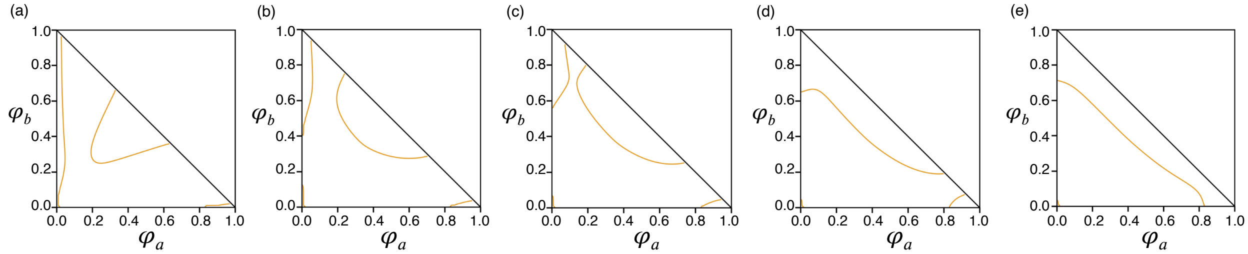


Figure S4: Phase diagrams showing the spinodal line (orange, solid) for a two component system with $u_{ab} = 5$ (a) $\Delta\epsilon_{bb} = \Delta\epsilon_{ab} = 0$, (b) $\Delta\epsilon_{bb} = \Delta\epsilon_{ab} = 0.8$, (c) $\Delta\epsilon_{bb} = \Delta\epsilon_{ab} = 0.9$, (d) $\Delta\epsilon_{bb} = \Delta\epsilon_{ab} = 1.0$ and (e) $\Delta\epsilon_{bb} = \Delta\epsilon_{ab} = 1.1$ . We set $N_a = N_b = 10, l = 2, z = 6, \bar{\epsilon}_{aa} = 1, \Delta\epsilon_{aa} = 0$, and $\bar{\epsilon}_{bb} = \bar{\epsilon}_{ab} = 0$ in all systems. See text *Section: Interrogating the effects of* $\Delta\epsilon_{ab}, \Delta\epsilon_{bb}$ *on the Phase Behavior of the Two Component System* for more details.

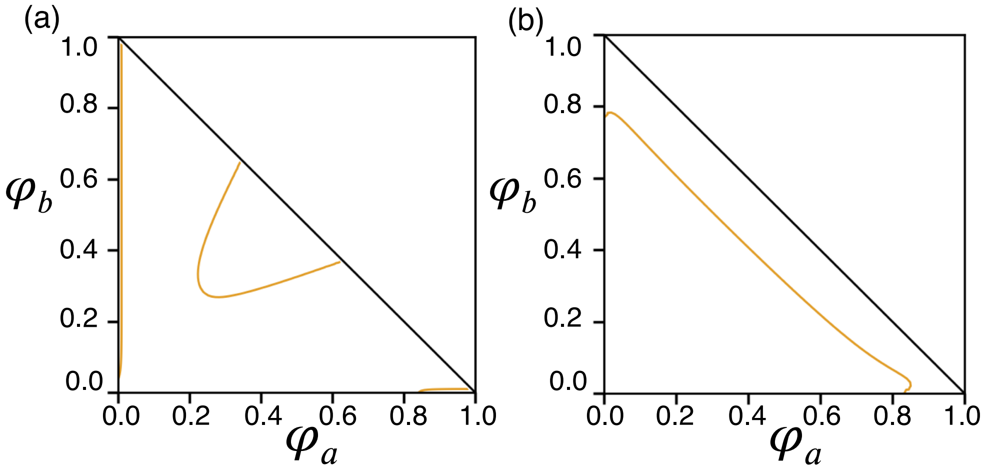


Figure S5: Phase diagrams showing the spinodal line (orange, solid) for a two component system with $u_{ab} = 5$ for a larger $N_a = N_b = 100$ system for (a) $\Delta\epsilon_{bb} = \Delta\epsilon_{ab} = 0$, (b) $\Delta\epsilon_{bb} = \Delta\epsilon_{ab} = 1.1$. Here we qualitatively recover the same phase behavior seen in the main text prior. We set $l = 2, z = 6, \bar{\epsilon}_{aa} = 1, \Delta\epsilon_{aa} = 0$, and $\bar{\epsilon}_{bb} = \bar{\epsilon}_{ab} = 0$ in all systems. See text *Section: Interrogating the effects of* $N, l$ *on the Phase Behavior of the Two Component System* for more details.

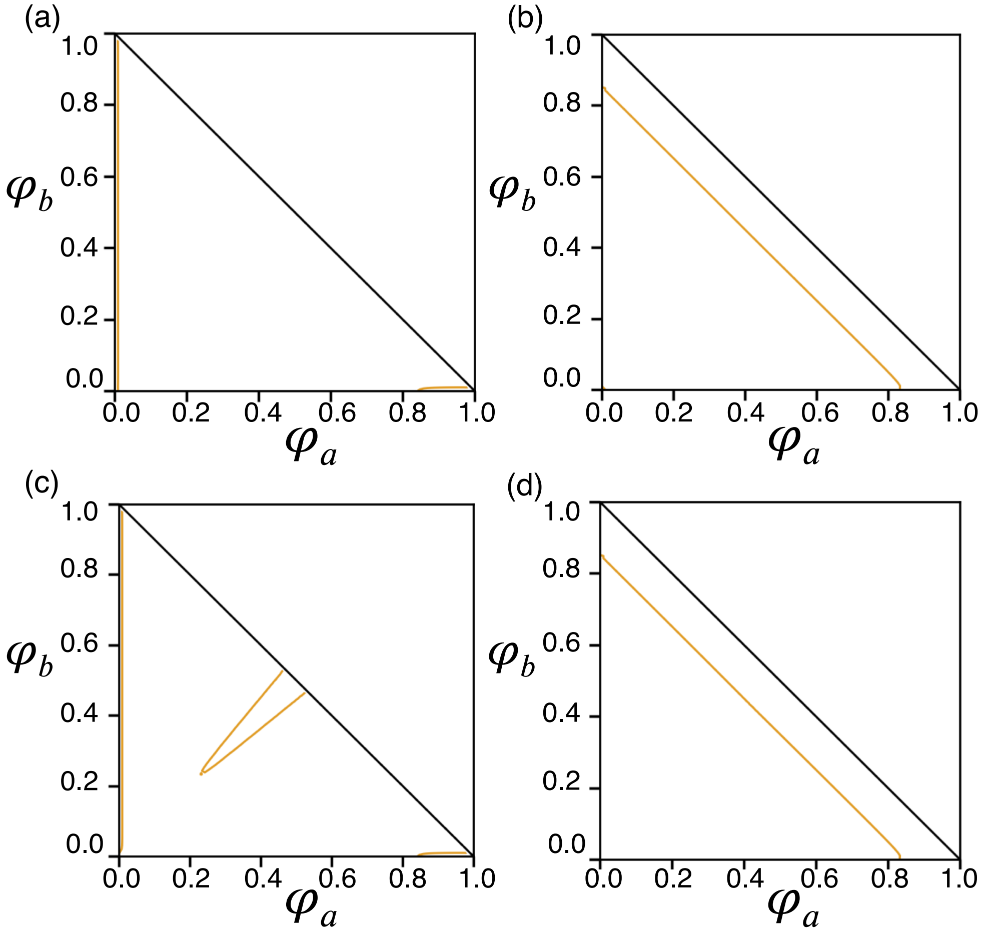


Figure S6: Phase diagrams showing the spinodal line (orange, solid) for a two component system with (a) $u_{ab} = 5, \Delta\epsilon_{bb} = \Delta\epsilon_{ab} = 0$, (b) $u_{ab} = 5, \Delta\epsilon_{bb} = \Delta\epsilon_{ab} = 1.5$, (c) $u_{ab} = 10, \Delta\epsilon_{bb} = \Delta\epsilon_{ab} = 0$, and (d) $u_{ab} = 10, \Delta\epsilon_{bb} = \Delta\epsilon_{ab} = 1.5$. We set $N_a = N_b = 100, l = 10, z = 6, \bar{\epsilon}_{aa} = 1, \Delta\epsilon_{aa} = 0$, and $\bar{\epsilon}_{bb} = \bar{\epsilon}_{ab} = 0$ in all systems. See text *Section: Interrogating the effects of $N, l$ on the Phase Behavior of the Two Component System* for more details.

(S1) Doi, M. *Introduction to Polymer Physics*, paperback ed.; Oxford University Press, 1990; p 136.

(S2) Rubinstein, M.; Dobrynin, A. V. Solutions of associative polymers. *Trends in Polymer Science* **1997**, *5*, 181–186.

(S3) Moré, J. J.; Garbow, B. S.; Hillstrom, K. E. *User Guide for Minpack-1*; Report, 1980.

(S4) Virtanen, P.; Gommers, R.; Oliphant, T. E.; Haberland, M.; Reddy, T.; Cournapeau, D.; Burovski, E.; Peterson, P.; Weckesser, W.; Bright, J.; van der Walt, S. J.; Brett, M.; Wilson, J.; Millman, K. J.; Mayorov, N.; Nelson, A. R. J.; Jones, E.; Kern, R.; Larson, E.; Carey, C. J.; Polat, İ.; Feng, Y.; Moore, E. W.; VanderPlas, J.; Laxalde, D.; Perktold, J.; Cimrman, R.; Henriksen, I.; Quintero, E. A.; Harris, C. R.; Archibald, A. M.; Ribeiro, A. H.; Pedregosa, F.; van Mulbregt, P.; SciPy 1.0 Contributors SciPy 1.0: Fundamental Algorithms for Scientific Computing in Python. *Nature Methods* **2020**, *17*, 261–272.

(S5) Jones, E.; Oliphant, T.; Peterson, P.; others SciPy: Open source scientific tools for Python. 2001–; `https://docs.scipy.org/doc/scipy/reference/generated/scipy.optimize.root.html#r9d4d7396324b-1`.

(S6) Sun, A. C.; Seider, W. D. Homotopy-continuation method for stability analysis in the global minimization of the Gibbs free energy. *Fluid Phase Equilibria* **1995**, *103*, 213–249.

(S7) Cots, O.; Gergaud, J.; Shcherbakova, N. *31st European Symposium on Computer Aided Process Engineering*; Elsevier, 2021; pp 1081–1086.

(S8) Shcherbakova, N.; Rodriguez-Donis, I.; Abildskov, J.; Gerbaud, V. A novel method for detecting and computing univolatility curves in ternary mixtures. *Chemical Engineering Science* **2017**, *173*, 21–36.

(S9) Shcherbakova, N.; Roger, K.; Gergaud, J.; Cots, O. *Computer Aided Chemical Engineering*; Elsevier, 2023; pp 709–714.

(S10) Bausa, J.; Marquardt, W. Quick and reliable phase stability test in VLLE flash calculations by homotopy continuation. *Computers & Chemical Engineering* **2000**, *24*, 2447–2456.

(S11) Bot, A.; Dewi, B. P. C.; Venema, P. Phase-Separating Binary Polymer Mixtures: The Degeneracy of the Virial Coefficients and Their Extraction from Phase Diagrams. *ACS Omega* **2021**, *6*, 7862–7878.

(S12) Allgower, E. L.; Georg, K. *Introduction to Numerical Continuation Methods*; Society for Industrial and Applied Mathematics, 2003.
